# Update on the mosquito fauna (Diptera: Culicidae) distribution in Cabo Verde: occurrence of the species complexes *Anopheles gambiae* and *Culex pipiens* (*pipiens*, *quinquefasciatus* and their hybrids)

**DOI:** 10.1101/2021.09.01.458512

**Authors:** Silvânia Da Veiga Leal, Isaias Baptista Fernandes Varela, Davidson Daniel Sousa Monteiro, Celivianne Marisia Ramos de Sousa, Maria da Luz Lima Mendonça, Adilson José De Pina, Aderitow Augusto Lopes Gonçalves, Hugo Costa Osório

## Abstract

In this study, we aimed to update the mosquito species composition and distribution based on a national entomological survey in all municipalities of Cabo Verde. This includes the sibling species of the *Culex pipiens* complex, namely *Cx. pipiens*, *Cx. quinquefasciatus* and their hybrids, in locations where information is not available. The entomological survey took place from October 2017 to September 2018, in all municipalities of Cabo Verde. Mosquito larvae and pupae were collected in breeding sites and samples were sent to the Laboratory of Medical Entomology of the National Institute of Public Health for the morphological identification of the species. The mosquitoes morphologically identified in *Anopheles gambiae* and *Culex pipiens* complexes of species were further molecular analysed to species confirmation. A total of 814 breeding sites were surveyed and 10 mosquito species of five genera were identified. The greatest number of mosquito species was reported in the island of Santiago. The most widespread species in the country were *Aedes aegypti* and *Culex quinquefasciatus. Anopheles arabiensis* was the only species identified in the *Gambiae* complex of species. The results of this study will assist decision makers in important health policies to control mosquitoes and vector-borne diseases towards a strategic approach by timely detection of changes in species diversity.

## Introduction

Vector-borne diseases represent a public health problem worldwide. At least 80% of the world population is at risk of infection by one or more diseases [1]. A large proportion of these diseases, such as malaria, dengue, chikungunya and Zika their infectious agents are transmitted by mosquitoes, which represent a major threat to human health, with millions of deaths annually [2,3].

In Cabo Verde some of the mosquito species, namely *Aedes aegypti* and *Anopheles arabiensis*, are vectors of pathogens, increasing the risk of diseases outbreaks. Moreover, the flow of people and goods between Cabo Verde and other countries poses a risk for the introduction of new mosquito species into the archipelago, as well as mosquito-borne pathogens. In the last 10 years, Cabo Verde has been affected by three mosquito-borne diseases outbreaks: dengue in 2009, Zika in 2015 and malaria in 2017 [3–7]. To evaluate the risk of transmission of vector-borne diseases in a given region, a continuous updating of the geographic distribution of insect vectors is required [8].

The first studies on the mosquito fauna in Cabo Verde date from the last century, after the identification of *Anopheles gambiae* s.l. in 1909 by Sant’Anna as described by Ribeiro and collaborators [9]. In 1977, an extensive study was carried out on all nine inhabited islands, resulting in the first and unique dichotomous key of the mosquitoes of Cabo Verde, in which eight species were described in the archipelago [10]. In 1984, the species *Lutzia tigripes* (formerly *Culex tigripes*) was first reported in the country [11]. The last update of Cabo Verde’s mosquito fauna was in 2007, based on bibliographic research, wherein entomological survey was carried out only in Maio, Santiago, Fogo and Brava islands. In that study, the species *Culex perexiguus*, a member of the *Culex univittatus* species complex was reported for the first time [12, 13]. Later, the species *Culex tritaeniorhynchus* was identified, raising to a total of 11 known species of mosquitoes recognized in the archipelago [14].

*Culex pipiens pipiens* and *Culex pipiens quinquefasciatus,* hereinafter referred as *Culex pipiens* and *Culex quinquefasciatus,* are vectors responsible for the transmission of lymphatic filariasis and neurotropic arboviruses to humans, namely West Nile virus [15,16]. These sibling species of the *Culex pipiens* complex were described in Cabo Verde in 1950 and 1980, respectively [10]. The nominal species of the complex, *Culex pipiens*, is found primarily in temperate zones, while *Cx. quinquefasciatus* occurs in warmer tropical and subtropical zones with a higher degree of humidity [17]. However, Cabo Verde is a region where both species coexist in sympatry and where hybrids of the two species were reported from Fogo and Maio locations [12,18]. Here, high levels of hybridization rates between the two species were detected, together with second-generation hybrids identified. The presence of these hybrid forms may locally potentiate the transmission of arboviruses to humans, such as West Nile virus, and therefore it is so important to have current information about its distribution.

In this study, we aimed to update the mosquito species composition and distribution based on a national entomological survey in all municipalities of Cabo Verde. This includes the sibling species of the *Culex pipiens* complex, namely *Cx. pipiens*, *Cx*. *quinquefasciatus* and their hybrids, in locations where information is not available. Due to the air and maritime transporting of people and goods in the archipelago, dispersion of species from one island to another, may have occurred after the last mosquito fauna survey in Cabo Verde in 2007. This could result in changes of composition, distribution and abundance of species, which potentially affects the pattern of disease occurrence and transmission. Our results will assist decision makers in important health policies to control mosquitoes and vector-borne diseases towards a strategic approach.

## Methodology

### Study area

Cabo Verde is a volcanic archipelago with an area of 4.033 km^2^ located about 550 km off the coast of Senegal in West Africa (Figure 1). The archipelago consists of ten islands, nine of which are inhabited with about 537.660 inhabitants. It has a warm and dry, arid and semi-arid climate, with an average annual temperature around 25°C, low rainfall, with two identified seasons: the dry season, from December to June, and the rainy season, from August to October [19].

**Figure 1:**
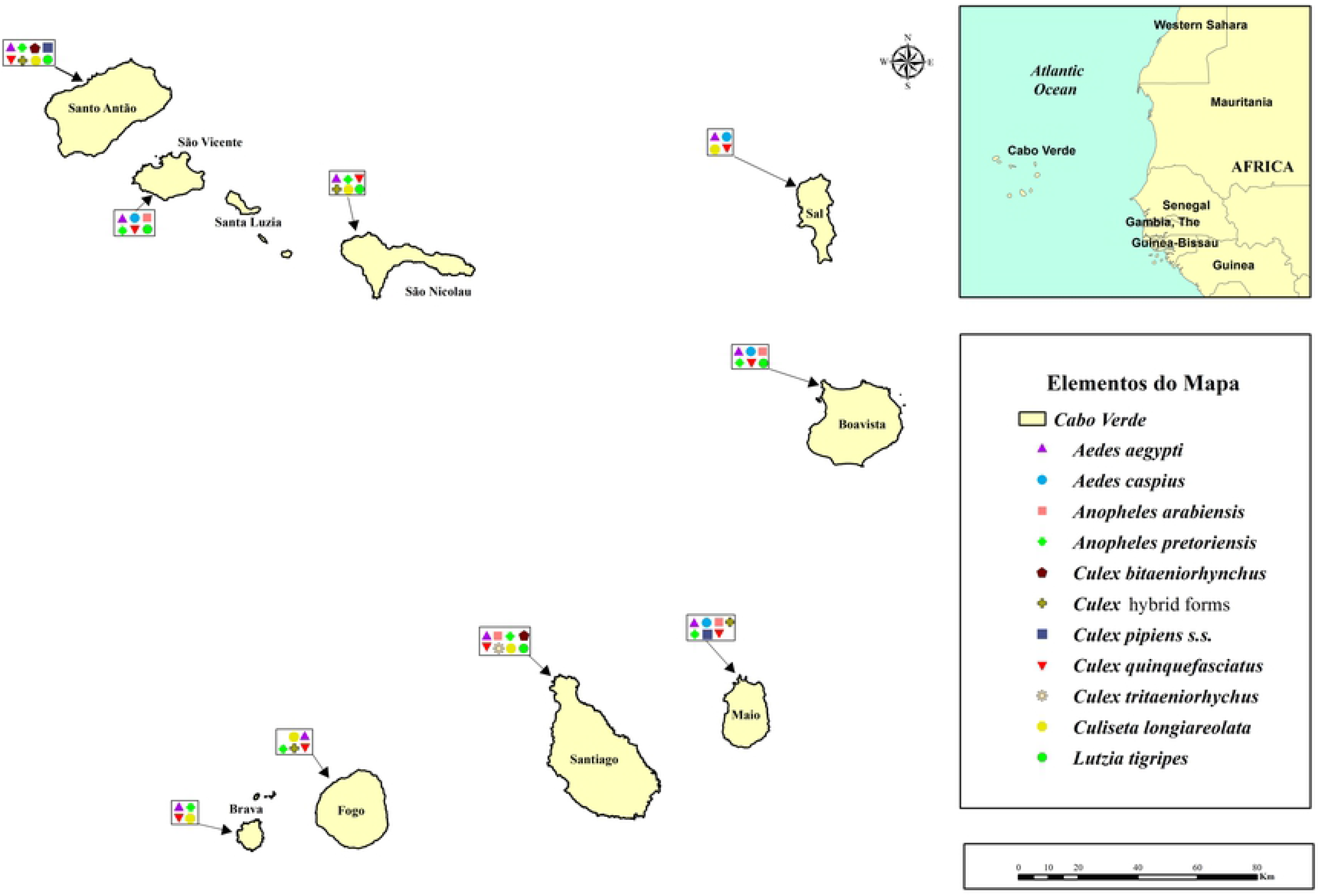
Geographic distribution of mosquito species in Cabo Verde (*ArcGIS* 10.5).

**Figure 2:**
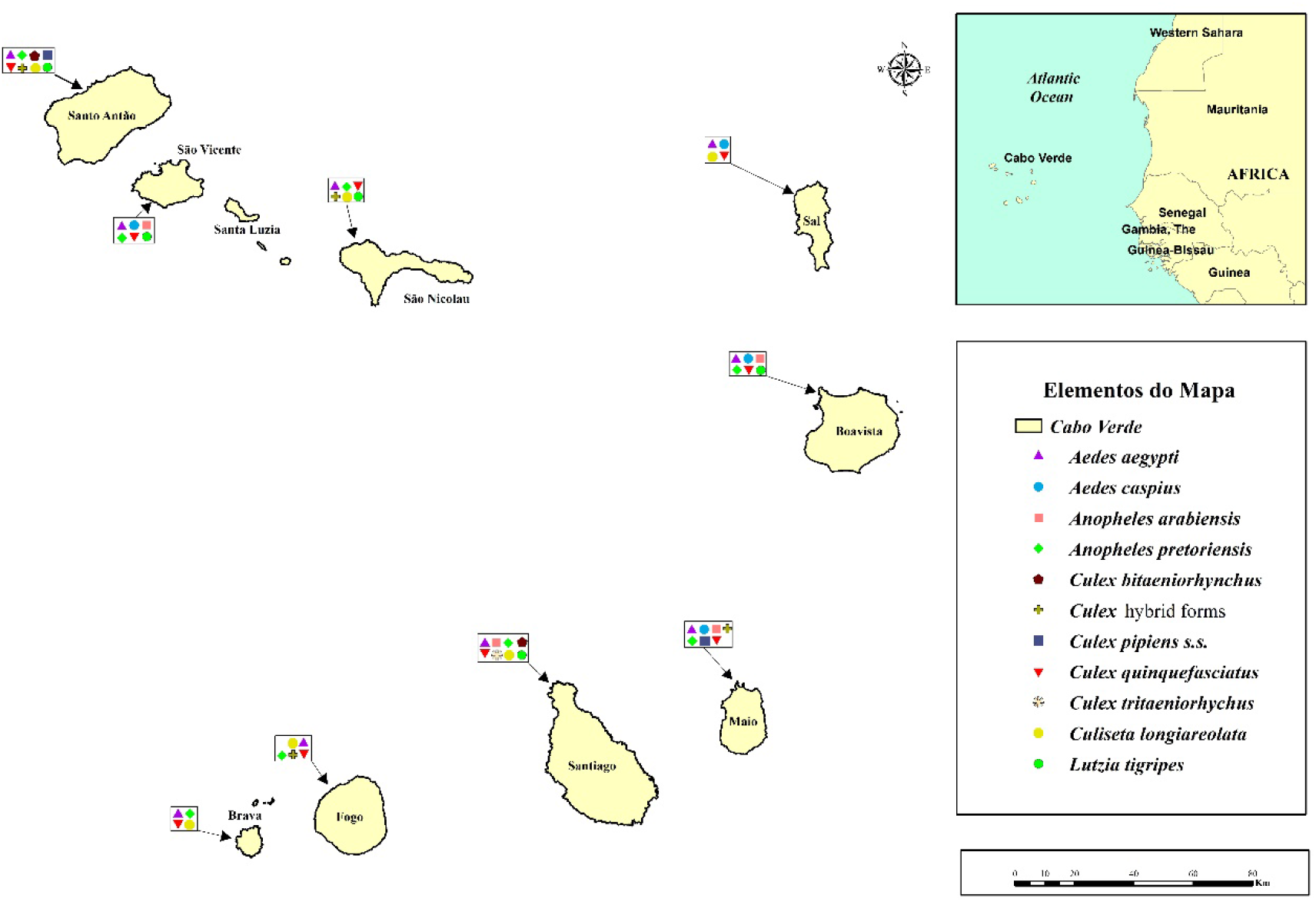
Geographic distribution of mosquito species in Cabo Verde (*ArcGIS* 10.5).

### Mosquito collection

The entomological survey took place from October 2017 to April 2018 in all municipalities of Cabo Verde, except in the island of Santo Antão, in which the survey took place in September 2018. Mosquito larvae and pupae were collected in breeding sites that included domestic containers and others. The samples were sent to the Laboratory of Medical Entomology (LME) of the National Institute of Public Health in 100 ml pots of water from the breeding site. Larvae L1 and L2 were kept in the insectary and fed with flocculated food for fish (Tropical mix flakes; Ref: F042.1), for later assembly of the L3/L4 stages in slides, prior species identification.

### Morphological identification

Larvae in development stage L3 and L4 and adults where used in the morphological identification of mosquito species. The larvae were mounted on slides with 2% glycerinated Hoyer’s medium. Morphological identification of the species was performed with the aid of a stereomicroscope using dichotomous keys according to Ribeiro et al. (1980) [10], Ribeiro & Ramos (1995) [9] and auxiliary taxonomic keys according to Dehghan et al. (2016) [20] and Azari-Hamidian & Harbach (2009) [21].

### Molecular analyses

The mosquitoes morphologically identified as *Anopheles gambiae* s.l. and *Culex pipiens* s.l. were submitted to single total DNA extraction using the NZY Tissue gDNA isolation kit. Sampled mosquitoes were individually grinded in Lysis Buffer using glass pearls in 2ml Eppendorf and DNA extraction was performed using the prepared lysate suspensions with the NZY Tissue gDNA isolation kit according to manufacturer’s recommendations. The Identification of *An. gambiae* complex species were performed by PCR-RFLP following the protocol described by Scott et al. (1993) [22] and Identification of *Cx. pipiens* species was performed according to Smith & Fonseca (2004) [23].

## Results

### Identification and geographical distribution of mosquito species

All the municipalities of the islands of Cabo Verde (N = 22) were inspected for mosquito larvae and pupae in a total of 814 breeding sites. Five genera and 10 mosquito species were identified (**Error! Reference source not found.**).

The greatest number of mosquito species was reported in the islands of Santiago and Santo Antão (N = 8); followed by Maio (N=7); Boavista, São Vicente, and São Nicolau (N = 6); Fogo (N=5), Sal and Brava (N = 4) (Figure 1).

The most widespread species in the country were *Aedes aegypti* and *Culex quinquefasciatus*, present in all islands of the archipelago (N= 09), followed by *Anopheles pretoriensis* (N = 08) and *Culiseta longiareolata* (N = 06). All the other species were circumscribed to less than five Islands. *Culex tritaeniorhychus* was the most restricted species found only on the island of Santiago, in the municipality of Tarrafal (Table 1).

**Table 1:**
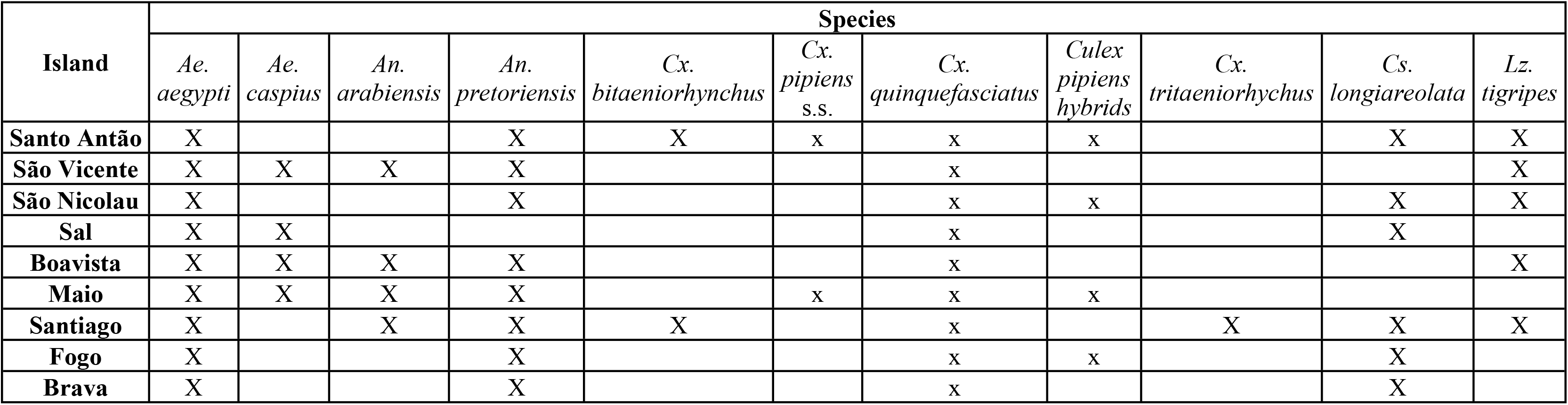
Species composition in the islands of Cabo Verde.

A total of 156 mosquitoes identified in the *An. gambiae* complex were molecularly identified as *An. arabiensis* (N = 156; 100%).

In the *Culex pipiens* complex the species the *Cx. quinquefasciatus* was the most identified species (N = 195; 83%) found in every island (Table 2). *Culex pipiens* was found in two islands, Maio and Santo Antão and in lower frequencies. Hybrid forms were identified in four islands (Maio, Fogo, Santo Antão and São Nicolau). Higher frequencies of hybrids were detected in Maio (N = 15; 50%). The species *Culex perexiguus* was not detected.

**Table 2.**
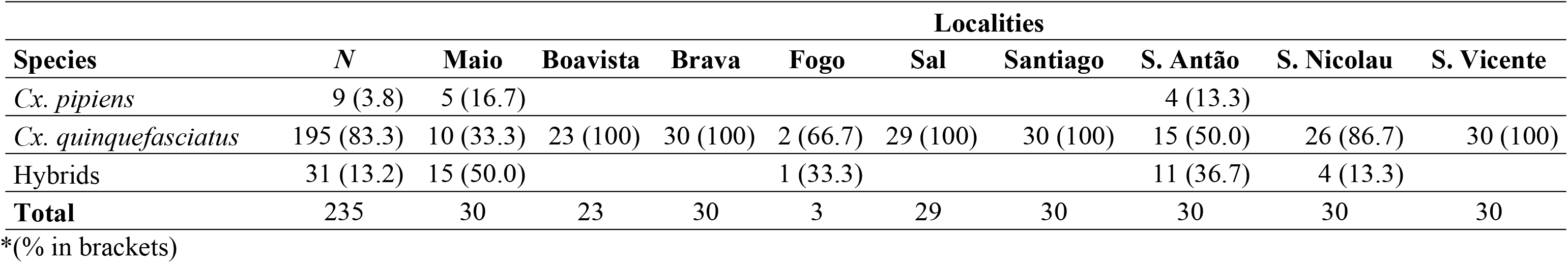
Frequencies of *Culex pipiens, Cx. quinquefasciatus* and their hybrids in Cabo Verde determined by the molecular assay ACE-2.

## Discussion

In this study, we present the most recent data on mosquito species distribution in Cabo Verde based on an entomological survey carried in all municipalities of the country. From the 10 species reported, four major mosquito vectors with medical importance were identified, namely *Anopheles arabiensis, Aedes aegypti, Culex pipiens* and *Cx. quinquefasciatus.*

The only malaria vector found in Cabo Verde was *An. arabiensis*, and it was identified for the first time in Maio and São Vicente in our survey. We corroborate previous studies reporting this species in Santiago, Fogo and Boavista [12, 24,25], however we found that its current distribution is wider. The exclusive presence of *An. arabiensis* in Cabo Verde can be explain by the Sahelian conditions of the archipelago, since this member of the *An. gambiae* complex is the most tolerant to aridity [26,27].

*Aedes aegypti* was identified in all 22 municipalities in the country. This was the second species of mosquito identified in Cabo Verde in 1930 in São Vicente. Half a century later, this species was identified on all inhabited islands except in Maio [10]. Although *Ae. aegypti* was not found in the island of Maio in the last entomological survey performed in 2007 [12], there was evidence of its presence, since the occurrence of cases of dengue and Zika in 2009 and 2015-2016, respectively [5,28]. In this survey, *Ae. aegypti* was reported in all the municipalities, and at municipality of Maio for the first time [29].

*Culex quinquefasciatus* and *Cx. pipiens* are the most ubiquitous mosquitoes in the tropical and temperate regions of the globe, respectively. In Cabo Verde, these species and their hybrids have been described since 1950 [10]. Entomological surveys have reported the presence of *Cx. pipiens* s.l. in all inhabited islands [10,12] which are corroborated in this study, in which we report its presence in all 22 municipalities. Although *Cx. pipiens* s.s. was identified in Maio and Santo Antão, which corroborates previous data [10], the presence of hybrid forms in Fogo and São Nicolau can suggest its presence in these islands. This brings up to four the locations of the archipelago that are more likely to present adequate environmental conditions for a contact zone between this species and *Cx. quinquefasciatus*.

*Culex quinquefasciatus* was identified in all inhabited islands of Cabo Verde. When in sympatry with *Cx. pipiens,* the former was always the most abundant form (N = 195; 83.3%).

Hybrid forms of *Cx. quinquefasciatus* and *Cx. pipiens* were found for the first time in Santo Antão. The subtropical location of the archipelago is likely to present adequate environmental conditions for a contact zone between the two species, as documented in previous studies [30]. Interestingly, there is no known hybrid zone between the two sibling species in mainland Africa, which probably reflects the effect of the Sahara Desert as a geographic barrier to the distribution of both species.

A recent study conducted on Santiago island showed a high human blood index (HBI) in engorged populations of *Cx. pipiens* s.l., followed by domestic dog and chicken blood meals (personal communication). Although depending on biotic and abiotic factors, including host-insect interaction factors, mosquito genetic traits, host availability and host density, it is generally assumed that each *taxa* has a particular biting behaviour, with *Culex pipiens* showing essentially an ornithophilic preference and *Culex quinquefasciatus* anthropophilic behaviour [31–43]. Hybrid forms can have intermediary host feeding patterns which can lead to a different role in pathogen transmission [35, 44–46].

*Aedes caspius* was identified in Sal, Boavista, Maio, and for the first time in São Vicente. *Aedes caspius* was found for the first time by Meira and collaborators in 1952 on the island of Sal [10,12].

Our results show *An. pretoriensis* widespread in Cabo Verde, except in Sal, where it was never reported (*Entomological Surveillance Bulletins*: https://insp.gov.cv/index.php/pilar-02-laboratorio-nacional-de-saude-publica, accessed in 27/03/2020). First detection of this species was in São Nicolau, in 1947. Later it was also reported in the islands of Boavista, São Vicente, Santo Antão, Santiago, Maio, Fogo and Brava [10].

*Culex bitaeniorhynchus* (formerly *Cx. ethiopicus*) was identified on the islands of Santo Antão and Santiago, where it has always been recorded, since its first identification, in 1977 [10,12]. And the *Culex tritaeniorhynchus* was found only on the island of Santiago in municipality of Tarrafal. This species was first described in the archipelago in 2011, in the municipality of Santa Cruz [14], which indicates dissemination of the species.

*Culiseta longiareolata* was recorded in Santo Antão, São Nicolau, Sal, Santiago, Fogo and Brava. This species was first described in 1947, in São Nicolau and Boavista, and in 1977 it was registered on all islands except Sal [8], while *Lutzia tigripes* was identified in Santo Antão, São Vicente, São Nicolau and Boavista. In Cabo Verde, this species was first reported in Maio [11], and later in Sal, Santiago and Fogo [12].

In this survey, the species *Culex perexiguus* was not detected. This species was reported for the first time in Santiago in 2010 [12], and since then no more reports of this species occurred. However, the confirmation of this species, included in the *Cx. univitattus* complex, in Cabo Verde is important, since this is a competent vector of West Nile, Sindbis, and Rift Valley fever viruses [47,48].

## Conclusion

Mosquitoes are the most important group of vectors in public health, because some species are highly competent for the transmission of pathogens to humans. The knowledge of the mosquito fauna and its geographic distribution provide important data to assess the potential transmission risk and outbreak occurrence.

With this work, 10 species of mosquitoes were identified in Cabo Verde, including members of two species complexes, four of which are important vectors of human pathogens, namely: *Anopheles arabiensis, Aedes aegypti, Culex pipiens and Cx. quinquefasciatus.*

The only malaria vector found in Cabo Verde was *An. arabiensis* and we report for the first time its presence in Maio and São Vicente. *Aedes aegypti* was spread across the archipelago and it was reported for the first time in Maio and *Aedes caspius* was reported for the first time in São Vicente.

Hybrids forms of *Cx. quinquesfasciatus* and *Cx. pipiens* were found in four islands, bringing up to four the locations of the archipelago that are more likely to these species occur in sympatry. In this survey *Cx. quinquefasciatus* was the most abundant species of the *Cx. pipiens* complex.

Timely detection of changes in species diversity provide valuable knowledge to health authorities, the scientific community and entities which may take control measures of vector populations reducing their impact on public health.

## Acknowledgements

We acknowledge the National Institute of Public Health of Cabo Verde for supporting the study, and the National Program for vector control to all the Health facilities. Also, we thank the health delegates, control vector agents, drivers, and administrators that provided their support during the entomological survey. We acknowledge Jonas Antonio Lopes Gomes, from the National Institute of Public Health of Cabo Verde, for the map drawing. Special acknowledge Tomás Alves de Só Valdez for trust and support.

## Authors’ contributions

Design of the study, S.D.V.L. and I.B.F.V.; Collections and laboratory work, D.D.S.M., C.M.R.d.S., A.A.L.G., and I.B.F.V.; Data analyses, I.B.F.V., A.A.L.G., H.C.O., and Original Draft Preparation, I.B.F.V., A.A.L.G., H.C.O., D.D.S.M., and S.D.V.L.; Writing-Review and Editing, I.B.F.V., A.A.L.G., M., A.J.D.P., H.C.O., and S.D.V.L.; Revision and Supervision, A.J.D.P., and M.d.L.L.M.

## Financing

The study was made possible through the Program Investing to achieve elimination for Malaria and impact against TB and HIV in Cape Verde, Grant CPZ-Z-CCSSIDA, supported by The Global Fund to Fight AIDS, Tuberculosis and Malaria and implemented by the Coordination Committee of the Fight against AIDS of Cape Verde (CCS-SIDA).

